# Berkson’s Paradox: A Common Statistical Error in Genomics Analyses

**DOI:** 10.64898/2026.10.22.761068

**Authors:** JJ Hum, Keegan Korthauer, Carl G. de Boer

## Abstract

Berkson’s paradox describes a scenario where a spurious correlation arises between two independent variables within a subset of samples selected based on a third variable which the first two variables cause. The conditions for Berkson’s paradox are perfect in genomics due to many variables we study being causally intertwined, common selection criteria being biased either intentionally or not by the variables of interest, and our propensity to look towards correlations for windows into underlying mechanism. Here, we provide examples of cases where spurious correlations are likely to arise in common genomics analyses, and suggest analysis approaches to avoid misinterpreting spurious correlations.

---

Tell me if you’ve heard this one before: An omics experiment is performed and, after data processing and comparison between samples, the extreme values are taken as a set. Next, this set is analyzed to learn something interesting about them. Other variables are considered to see if there are correlations between them that may suggest why this set is special. A negative association is found between variables X and Y. This is very exciting because, for most genes, X and Y are positively correlated! Perhaps this gene set is unique and is regulated in a special way?

This is a common scenario in omics analyses. While the details can take many forms, the core remains the same: selecting a subset by skimming the extremes of a distribution and then considering the relationships between variables within this set; variables that often are related to the original selection criteria. The biased selection process can result in Berkson’s paradox, resulting in misleading correlations between variables^1^. The spurious correlations can be positive, negative, or even a false lack of correlation; it depends on the relationships between the variables and the nature of the selection.

Berkson’s paradox is caused by the biased selection process inducing a relationship between the two variables being compared. It is perhaps easiest to illustrate with an example (Fig. 1): Say two transcription factors (TFs), TF_A_ and TF_B_, bind DNA and make chromatin more accessible. In order to be one of the top most accessible regions, there must be a critical amount of binding for the sum of TF_A_ and TF_B_. At one extreme, the threshold is passed solely due to TF_A_ and no TF_B_ binding is needed. At the other, there is no TF_A_ and the only way to pass the threshold is with enough TF_B_. In this biased subset of the data, we have induced a quantitative relationship between TF_A_ and TF_B_: where there is more TF_A_, there is less TF_B_. This resulting negative correlation between binding of TF_A_ and TF_B_ has nothing to do with whether the two influence each other’s binding. Instead, it results from the selection process being related to the variables being considered.

**Figure 1:**
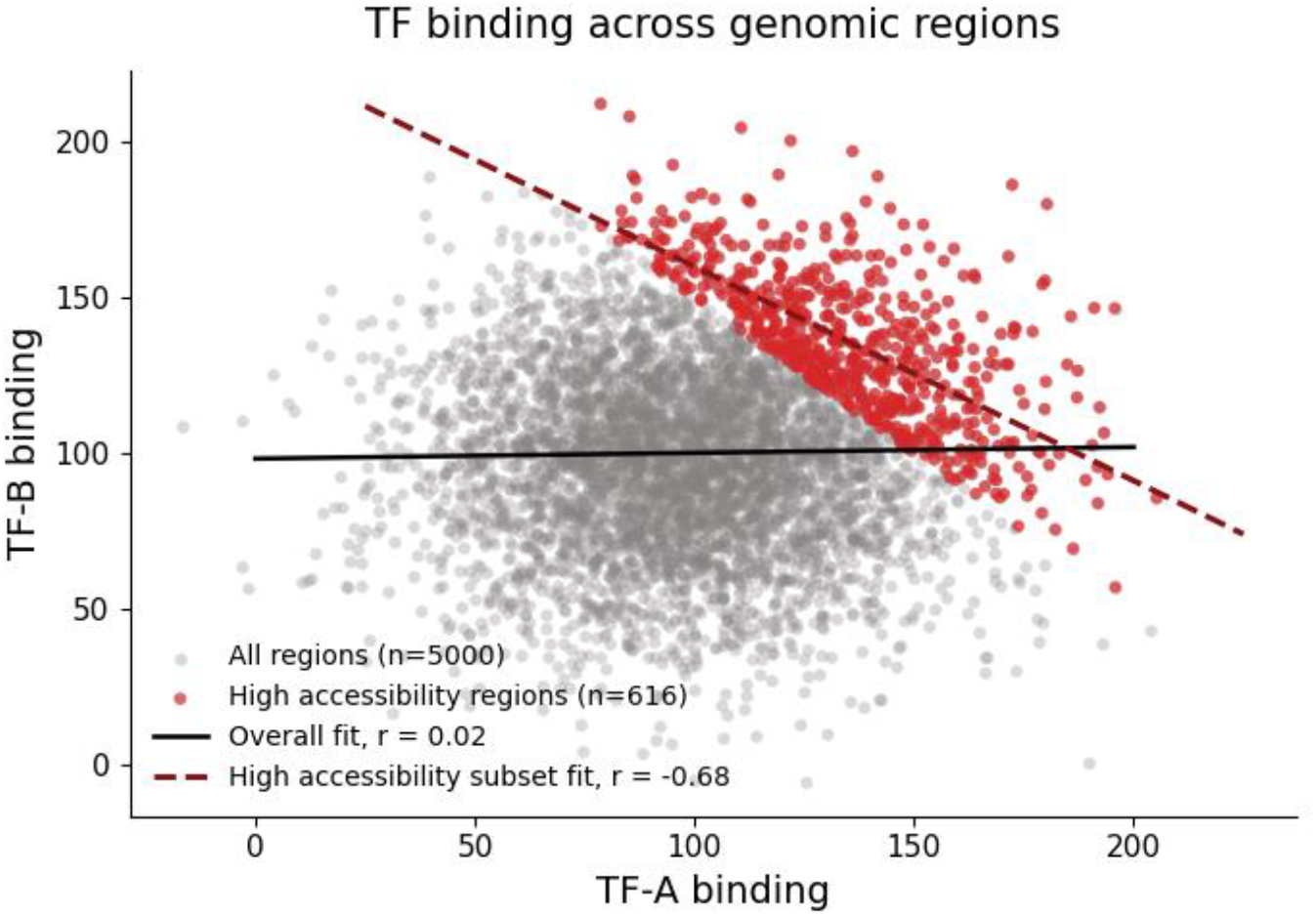
Illustration of Berkson’s paradox. A scatterplot showing binding for two hypothetical TFs (*x-* and *y*-axes). The binding of the TFs is uncorrelated across all datapoints (gray points, solid black line of best fit), but if both TFs independently influence accessibility and we select for only high accessibility regions (red), their binding becomes anticorrelated (dashed red line of best fit).

Berkson’s paradox can also happen when considering the same variable in different ways. For instance, if we inspect ChIP-seq data for a TF, we might find that, within the regions with the highest ChIP signal, the TF’s affinities for each DNA sequence are spuriously anticorrelated with its number of binding sites. This is because being a top ChIP-seq peak requires some combination of many low-affinity sites or few high-affinity sites. Again, this negative correlation is expected even if there is no relationship between binding site affinity and number. If we performed ChIP-seq of completely random DNA, where motifs for the TF occurred purely by chance, a negative correlation between binding site number and affinity would be expected because they both cause the ChIP-seq signal.

Berkson’s paradox is a type of collider bias, where a relationship between two variables is altered by consideration of a third (in this case the selection criteria)^2^. In some cases, the two variables had some relationship that is now perturbed (e.g. negated, reversed). In other cases, the two variables had no relationship to each other, and now have a relationship (either positive or negative) by virtue of them both having relationships to the selection criteria.

Many selection approaches used in genomics can lead to Berkson’s paradox. All it requires is a biased selection process for what entities (e.g. peaks, genes, cells, individuals) are being considered, and the variables considered being related to the selection process. The selection bias can result from intentionally biased selection or even statistical power that favor extremes of the distribution, both of which are common in genomics. Most biochemical phenomena we are measuring represent a continuum, where we can detect smaller and smaller effects as the data improve (e.g. via more replicates, sequencing depth). For instance, ChIP-seq peaks and open chromatin regions are defined statistically, requiring some enrichment relative to a background model. Stronger signals are more statistically robust, and so more likely to end up selected. This means that we are inherently skimming the top of the distribution^3^. Imagine we had done only two replicates of ATAC-seq to measure open chromatin, both of which we under-sequenced. The only accessible regions we can confidently identify are the extreme cases. Amongst these, it is now expected that, for chromatin-opening TFs, their binding levels are anticorrelated because their activities were interchangeable (Fig. 1). Now imagine that two identical studies were performed, one deeply sequenced and capturing much of the distribution, and the other shallowly sequenced, sampling only the top of the distribution. In this case, the open chromatin regions identified in these two studies could show conflicting correlation patterns between TFs binding in open chromatin, not because there are true differences, but because one was sequenced deeper.

Biased selection is common in differential analysis. The statistical significance of a difference between conditions (e.g. RNA expression changes between treatment and control) depends on how different the conditions are, the number of samples, and the variance of the data. Increasingly weak differences will be detectable (i.e. statistically significant) with better experimental design (e.g. more replicates, sequencing depth)^4^. Consequently, as before, our selection criterion (e.g. significantly differentially expressed genes) gets more extreme as data quality lowers. However, differential analysis can be even more problematic because a threshold is often explicitly applied (e.g. minimum log fold change) with the intent of skimming the most extreme values from the distribution, resulting in an intentionally biased sampling of entities for study.

Similar issues can happen in single cell analysis. For instance, it is common to find interesting cell states (cell types, or other interesting subsets) using gene expression signatures (e.g. a weighted sum of gene expression values)^5^. In this case, genes that mark the cell state would likely be strongly positively correlated across all cells, expressing highly in the cell state of interest and lowly in others. Within the cell state, however the signature genes may be anticorrelated because they were interchangeable in meeting the selection criteria.

In genomics, our variables of interest are often related to our selection criteria, potentially leading to collider bias. Many of the biochemical activities associated with gene regulation are correlated with each other, including TF binding, open chromatin, histone marks, DNA methylation, and gene expression. There are many feedback loops in gene regulation, and each of these biochemical activities is thought to cause the others^6–8^. For instance, TF binding can change DNA methylation and DNA methylation can change TF binding^9–11^; transcription can change histone marks and histone marks can change transcription^12–14^. Consequently, selection based on any of these could cause misleading correlations between the others. To illustrate this, we determined how correlations between histone mark ChIP-seq signals would change when considering all promoter regions versus promoters of the most highly expressed genes (Fig. 2).

**Figure 2.**
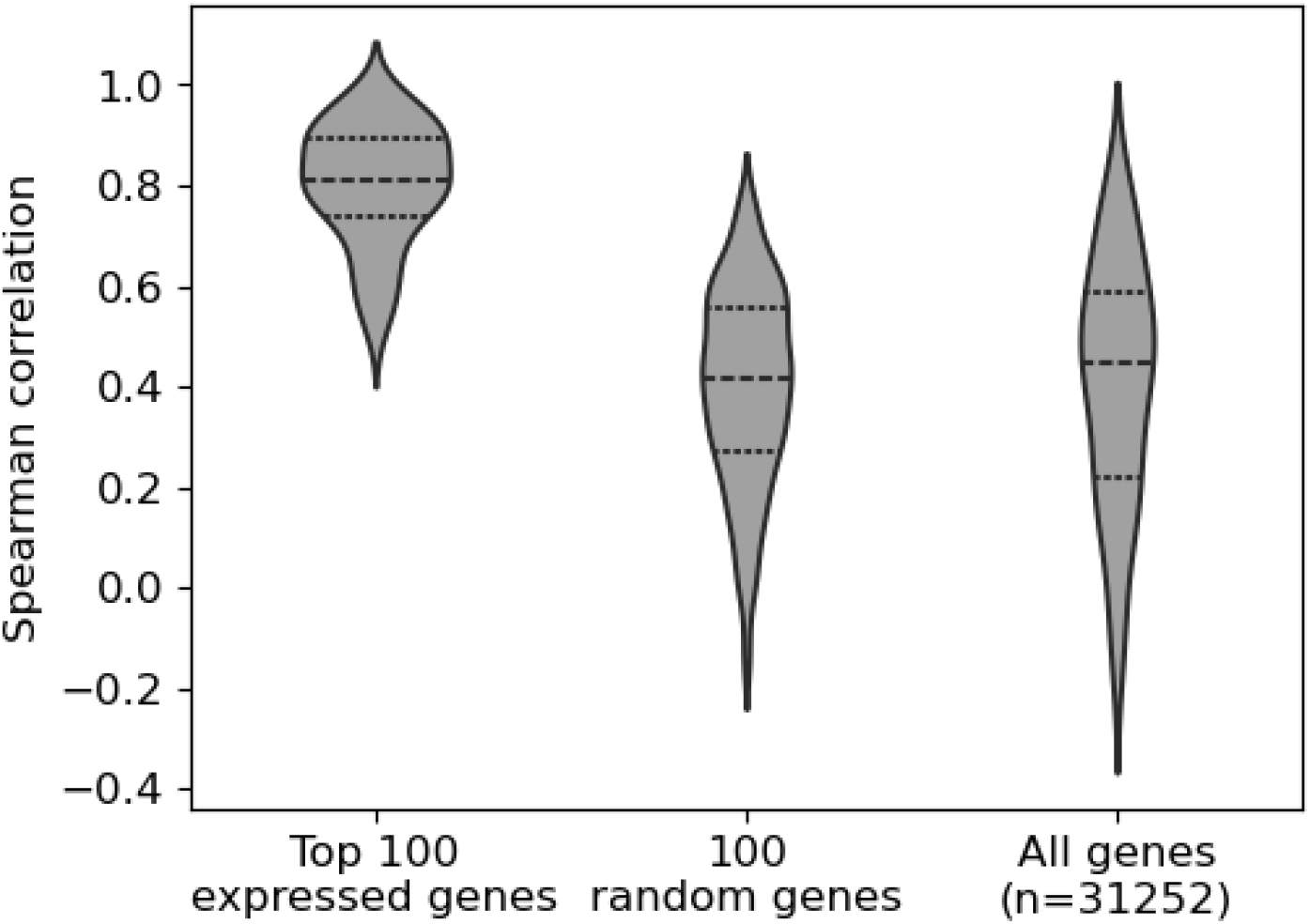
Sampling only the top expressed genes produces distinct correlations between histone ChIP-seq signals in promoter regions. Spearman correlation of histone ChIP-seq reads in promoter regions (defined as +/-2500 bp around the transcription start site; y-axis) across the top 100 highly expressed genes, 100 random genes, and all genes (x-axis). Dotted lines represent the top and bottom quartiles, with dashed midline indicating the median.

One area of genomics where Berkson’s paradox is well known is statistical genetics^1^. In a population with a heritable disease, the frequencies of disease-associated SNPs may appear to be anticorrelated. This is because a threshold genetic liability is required to get disease. For example, in a scenario where you require any one of variant A or variant B to get a disease, most individuals with the disease will have A but not B, or B but not A. Rarely, they might have both, but they never have none otherwise they would not have the disease. This leads to a negative correlation between the frequencies of A and B in the diseased population. Further, if A impacts disease progression (e.g. severity, mortality), this will induce a spurious negative correlation between B and disease progression. Several statistical methods have been developed to mitigate this bias. For instance, Mahmoud et al. use clustering of SNP effect sizes to identify variants that affect only disease incidence and not progression, and derive correction factors based on those^15^. Alternatively, a pseudo-population that represents the true underlying population can be created by using inverse probability weighting to scale selected observations by the inverse of their probability of being selected^16^.

Given that many common genomics analyses are likely to lead to Berkson’s paradox, it is surprisingly poorly known in the genomics community. Real damage can arise from the incorrect conclusions drawn from it as people work to nail down a mechanism suggested by a spurious correlation that runs counter to reality. Making Berkson’s paradox and collider bias an integral part of a standard genomics education would help prevent misapplication and overinterpretation of correlations.

## Methods

For Figure 1, TF-A and TF-B values were sampled from a normal distribution (μ = 100; σ = 30). Accessibility was calculated by adding TF-A and TF-B values, and a high accessibility subset was created by selecting points where accessibility was greater than an arbitrary threshold of 250. Pearson’s correlations between TF-A and TF-B were calculated for the subset and full set of values.

For Figure 2, K562 RNA-seq data was obtained from ENCODE (experiment ENCSR792OIJ, file ENCFF928NYA). The promoters for each gene were extracted from the GENCODE V49lift37 (knownGene) table and corresponded to the regions surrounding each gene’s transcription start site (+/-2500 bp). Three promoter sets were created and analyzed for correlations between histone marks: all promoters, those for the top 100 expressed genes, and those for 10 replicates of 100 randomly selected genes (to control for effects that are simply due to a decreased gene set size). K562 histone ChIP-seq peak data for 12 different histone marks was obtained from ENCODE (pooled reads from 2-3 biological replicates; Table 1). The ChIP signal for each promoter region was extracted for each histone mark using deepTools multiBigwigSummary, then Spearman correlations were calculated for each histone pair (66 unique pairs) across promoter regions for each promoter set using deepTools plotCorrelation^17^.

**Table 1.** K562 histone ChIP-seq data obtained from ENCODE.

| ENCODE Experiment | ENCODE Filename | Target |
| --- | --- | --- |
| ENCSR000AKX | ENCFF694ODT | H4K20me1 |
| ENCSR000AKR | ENCFF605EVL | H3K36me3 |
| ENCSR000AKQ | ENCFF582IMB | H3K27me3 |
| ENCSR000APD | ENCFF334HSS | H3K79me2 |
| ENCSR000AKW | ENCFF220RGS | H3K9me1 |
| ENCSR000AKT | ENCFF054RSU | H3K4me2 |
| ENCSR000AKV | ENCFF239EBH | H3K9ac |
| ENCSR000APC | ENCFF202EVH | H2AFZ |
| ENCSR000AKP | ENCFF465GBD | H3K27ac |
| ENCSR000AKS | ENCFF457URZ | H3K4me1 |
| ENCSR000APE | ENCFF632NQA | H3K9me3 |
| ENCSR668LDD | ENCFF405ZDL | H3K4me3 |

